# Spatial Multi-omics of Arterial Regions from Cardiac Allograft Vasculopathy Rejected Grafts

**DOI:** 10.1101/2023.04.12.536667

**Authors:** Jessica Nevarez-Mejia, Harry C. Pickering, Rebecca A. Sosa, Gregory A. Fishbein, William M. Baldwin, Robert L. Fairchild, Elaine F. Reed

## Abstract

**Background:** Cardiac allograft vasculopathy (CAV) is a major cause of late-graft failure and mortality following heart transplantation. The mechanisms underlying vascular remodeling are poorly understood. A major immune risk factor associated with the development of CAV is the presence of donor-specific antibodies (DSA) that induce chronic endothelial cell (EC) injury, leukocyte recruitment and inflammation resulting in thickening of arterial intima. Here, we molecularly characterized innate and adaptive immune cells present in the arteries of rejected cardiac allografts with DSA and identified protein and transcriptomic signatures distinguishing early and late CAV lesions.

**Methods:** Arterial areas of interest (AOIs) from CAV+DSA+ rejected cardiac allografts (N=3; 2 females, 1 male) were subjected to GeoMx digital spatial profiling (DSP). AOIs were scored on the level of CAV progression/neointimal thickening (22 AOIs total; 11 high and 11 low neointima) and were subjected to whole transcriptome and protein profiling.

**Results:** AOIs with low neointima significantly increased markers for activated inflammatory infiltrates, transcripts of EC activation and gene modules involved in activating metalloproteinases and TP53 regulation of caspases. Inflammatory and apoptotic protein markers significantly correlated with inflammatory modules in AOIs with low neointima. AOIs with high neointima increased TGFβ-regulated transcripts and modules enriched for platelet activation/aggregation. Proteins encoding SMCs and growth factors/survival correlated with modules enriched for proliferation/repair in AOIs with high neointima. Key transcripts in promoting proliferation, migration, and EndoMT were significantly associated with increasing neointima scores.

**Conclusion:** Our results reveal new protein and transcriptomic signatures associated with CAV progression. Lesions exhibiting inflammatory profiles appear to be early lesions that transition to later proliferative/pro-fibrotic phenotype CAV lesions. These findings should form the foundation for the identification of improved biomarkers to guide CAV treatment.

## 3. Introduction

Cardiac allograft vasculopathy (CAV) remains a major clinical challenge limiting long-term graft and patient survival following heart transplantation. Approximately 29% of heart transplant recipients develop CAV by 5 years and 47% by 10 years post-transplant^1^. CAV lesions are characterized by concentric intimal thickening of the vascular wall consisting of proliferating myofibroblast and inflammatory immune infiltrates^2^. Although both immunological and non-immunological processes contribute to CAV pathogenesis, the exact mechanisms mediating disease progression remain unclear. Episodes of antibody-mediated rejection (AMR) in which donor specific antibodies (DSA) target human leukocyte antigens (HLA) present on vascular endothelial cells (ECs) have been increasingly recognized as a major risk factor contributing to CAV^3^. Specifically, DSA can activate EC intracellular signaling cascades inducing EC proliferation, migration, and increased surface expression of adhesion molecules promoting monocyte and NK cell recruitment^4–8^. Moreover, DSA can directly mediate EC-injury by triggering activation of the classical complement cascade^8^. Recurring episodes of AMR contribute to chronic inflammation eliciting EC injury and apoptosis. CD4+ and CD8+ T-cells activated by HLA alloantigen recognition are also commonly found in the adventitia and neointima of CAV rejected hearts. Specifically, memory T helper (Th) 1 secreting pro-inflammatory cytokines such as IFN-у and TNF-α can attract and activate CD8+ cytotoxic T cells and NK cells^2^. T cells also secrete transforming-growth factor-β (TGF-β) which significantly contributes to fibrosis and remodeling by upregulating collagen synthesis in SMCs and stimulating macrophage secretion of matrix metalloproteinases^9^. The process of endothelial-to-mesenchymal transition (EndoMT) remains another distinguishing characteristic contributing to CAV. EndoMT is a phenomenon in which ECs gradually lose EC markers and increase mesenchymal cell. A combination of inflammatory stimuli, low or disturbed shear flow, vascular stiffness (activating WNT/β-catenin), and metabolic dysregulation (e.g., high glucose) can all cause EndoMT via different signaling pathways^10^.

Overall, numerous immunological mechanisms involving both innate and adaptive immune responses have been shown to contribute to the initiation and progression of neointimal development in CAV. More recent studies using endomyocardial biopsies (EMBs) have elucidated DSA and/or AMR-specific bulk-RNA signatures which contribute to CAV^11–13^. However, as CAV mainly affects large and small epicardial and intramyocardial arteries, EMBs (taken from the right ventricle) may not exactly reflect the immune and/or vascular signatures contributing to neointima formation. Therefore, the proteomic and transcriptomic signatures revealing specific activated markers and regulatory pathways from DSA+CAV+ affected vessels as observed in rejected cardiac explants remains to be elucidated. In this study, we used GeoMx digital spatial profiling (DSP) to examine both the whole transcriptome and targeted protein profiles of arterial regions scored with varying levels of neointima progression. Using a multi-omics approach, we explored the differences between arterial regions containing low and high neointimal scores thereby identifying potential immune proteins, enriched transcripts, and pathways involved in neointima progression.

## 4. Methods

### 4.1. Study approval

The use of human cardiac rejected explants for this study was approved by the University of California, Los Angeles Institutional Review Board (IRB#18-001275). Formalin-fixed paraffin embedded (FFPE) tissue from rejected cardiac explants were obtained from the UCLA Translational Pathology Core Lab (TPCL).

### 4.2. Arterial vessel selection and pathological characteristics

A total of 22 arterial regions were selected across three cardiac allograft explants from CAV+DSA+ patients (N=3; 2 females PID1, PID2 and 1 male PID3). Patient demographics and clinical information are summarized in **Supplementary Table 1**. These arterial regions included 10 vessels from PID1, five vessels from PID2, and seven vessels from PID3. Hematoxylin and eosin (H&E) staining was used to score arteries based on the level of CAV progression/neointimal thickening by the pathologist. In total, 11 arteries were scored with ‘low’ neointima (+/- minimal and 1+ mild) and 11 with ‘high’ neointima (2+ moderate, 3+ significant, and 4+ very significant) (**Supplementary Table 2**). H&E images were captured using Zeiss microscope and examined using ZEISS ZEN lite Software v3.3.

### 4.3. Targeted protein and whole transcriptome GeoMx digital spatial profiling (DSP)

Multiplex digital spatial profiling of protein and RNA in fixed tissues was performed by NanoString Technologies (Seattle, WA) as part of the Technology Access Program (TAP) as per Merritt et al., *Nat Biotechnol* 2020^14^ (**Supplementary Methods 1.1**).

### 4.4. Data normalization using NanoString’s GeoMx analysis suite

The GeoMx analysis suite was used for quality control (QC) and normalization of both protein and RNA datasets (**Supplementary Methods 1.2**).

### 4.5. WGCNA analysis, gene regulatory networks (GNRs), and data deconvolution

Whole gene co-expression network analysis (WGCNA) was used to define modules of co-expressed transcripts^15^. Gene regulatory network (GRN) analysis was performed using R package *SCENIC*^16^. RNA deconvolution was derived using SpatialDecon algorithm as previously described^17^ (**Supplementary Methods 1.3**)

### 4.6. Multiplex-immunofluorescent staining

Multiplex-immunofluorescent staining of cardiac explants was performed by UCLA TPCL (**Supplementary methods 1.4**).

### 4.7. Statistical Analysis

Identification of differentially expressed proteins (DEPs), differentially expressed genes (DEGs), and differentially expressed WGCNA modules between AOIs with low and high neointima were determined using linear mixed-effects model, including patient ID as a random effect variable. Results were considered significant at Log_2_FC > 1 and *p-value* < 0.05 for **Figures 3B, C** (**Table. 1**) and *p-value* < 0.05 for **Figure 3D**. Statistical significance in **Figure 2C** was determined using Pearson correlation coefficient test, significant at *p-value* < 0.05. Results in **Table 2** (and **supplementary Figure 4**) were determined using Spearman correlation *p* < 0.05 (*), *p* < 0.01(**), *p* < 0.001(***) in GraphPad Prism software v9.3.1.

**Table 1.** Differentially expressed proteins (DEPs) and differentially expressed genes (DEGs) between low and high neointima AOIs.

|  | **Protein** | **Log<sub>2</sub>FC** | ***p*-value** |
| --- | --- | --- | --- |
| **1** | OX40L | 2.82 | *0.029* |
| **2** | CD45RO | 1.23 | *0.019* |
| **3** | pan-RAS | 1.22 | *0.027* |
| **4** | CD14 | 1.17 | *0.035* |
| **5** | INPP4B | 1.15 | *0.049* |
| **6** | CD127 | 1.14 | *0.023* |
| **7** | p53 | 1.12 | *0.046* |
| **8** | Tim-3 | 1.11 | *0.041* |
|  | **Gene** | **Log<sub>2</sub>FC** | ***p*-value** |
| **1** | *VWF* | 1.42 | *0.021* |
| **2** | *CSRP1* | -1.05 | *0.022* |
| **3** | *TAGLN* | -1.18 | *0.012* |
| **4** | *ADIRF* | -1.33 | *0.009* |

**Table 2.**
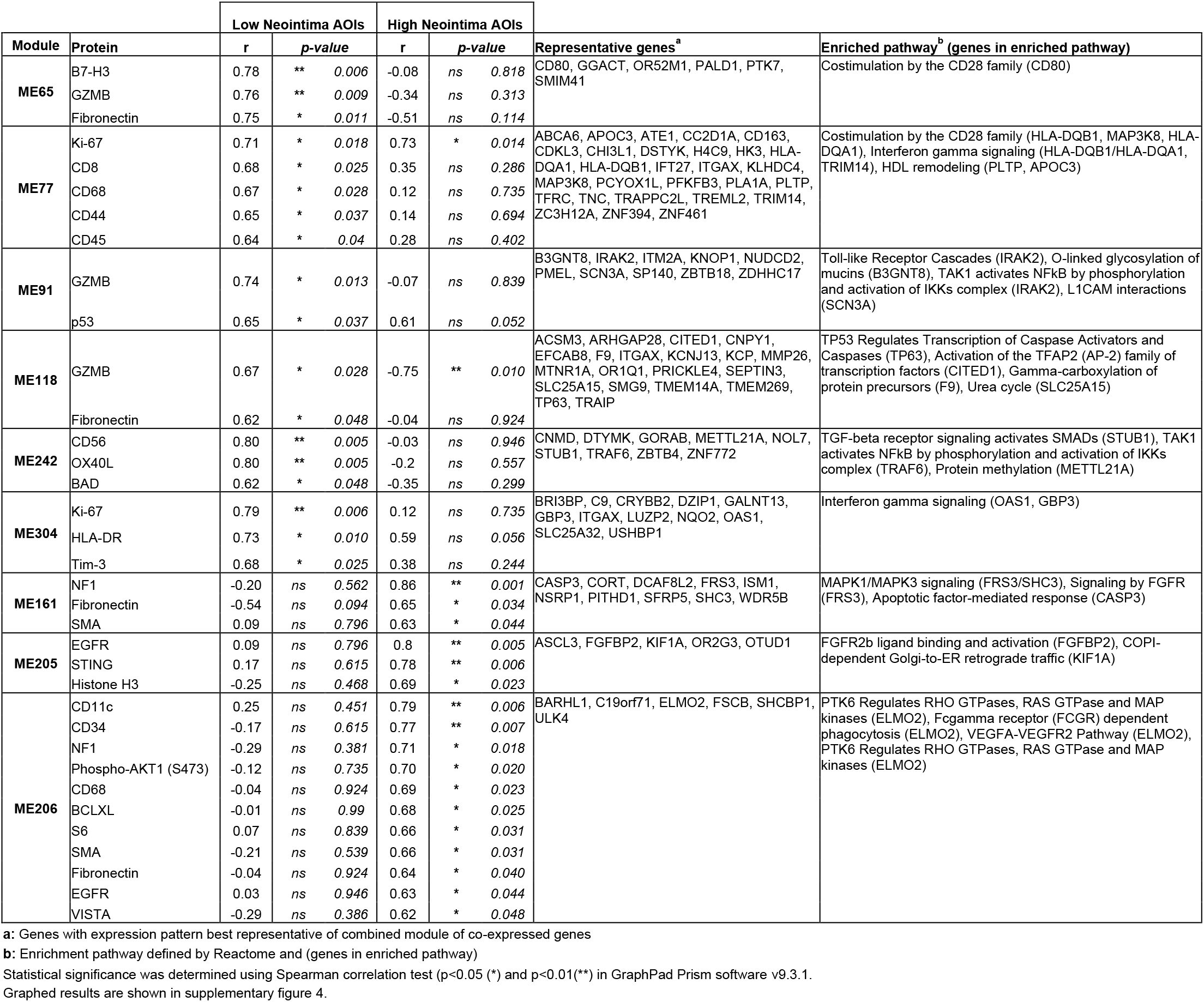
Highly correlated proteins with WGCNA modules between low and high neointima AOIs.

## 5. Results

### 5.1. Arterial AOIs express protein markers related to immune-cell activation and cell death

Arterial AOIs from three CAV+DSA+ rejected cardiac explants (PID1-3; 2 females and 1 male; **Figure 1A**) were spatially profiled using a 73-protein panel containing markers relating to immune activation, cell death, and signaling pathways (**Supplementary Table 3**). A total of 41 protein markers were similarly expressed across all arterial AOIs (SNR>1 in blue) (**Figure 1B**). Mesenchymal markers, SMA and Fibronectin were highly expressed, indicative of vessel region selection. Markers involved in immune cell activation and cytotoxicity (CD44, GZMB and HLA-DR), apoptosis (cleaved caspase 9, BAD, and p53) and cell survival (BCLXL) showed moderate expression. This was accompanied by the expression of CD45+ immune infiltrates (median SNR 10.56) including macrophages (CD68), T cells (CD4 and CD8), NK cells (CD56), monocytes (CD14), and dendritic cells (CD11c) in order of expression. Phosphorylated proteins in the MAPK and PI3K/AKT signaling pathway (p44/42 MAPK ERK1/2, pan-AKT, and phospho-AKT1) were detected at moderate-to-low levels. Stimulator of interferons genes (STING) and TNFSF4/OX40L, a memory T cell survival inducing also showed moderate-to-low levels of expression (**Supplementary Table 4**). Protein correlation matrix analysis across all AOIs revealed significant associations between apoptotic and anti-apoptotic proteins (e.g., BAD and BCLXL), T cell markers (CD3 and CD8), and signaling cascades (Phospho-AKT1 (S473) and Pan-Ras) (**Supplementary Figure 1 and Supplementary Table 5**). Overall, the expression of these markers suggests that arteries from rejected cardiac explants experience high levels of vascular inflammation encompassed by leukocyte infiltration, cytotoxicity, and apoptosis.

**Figure 1.**
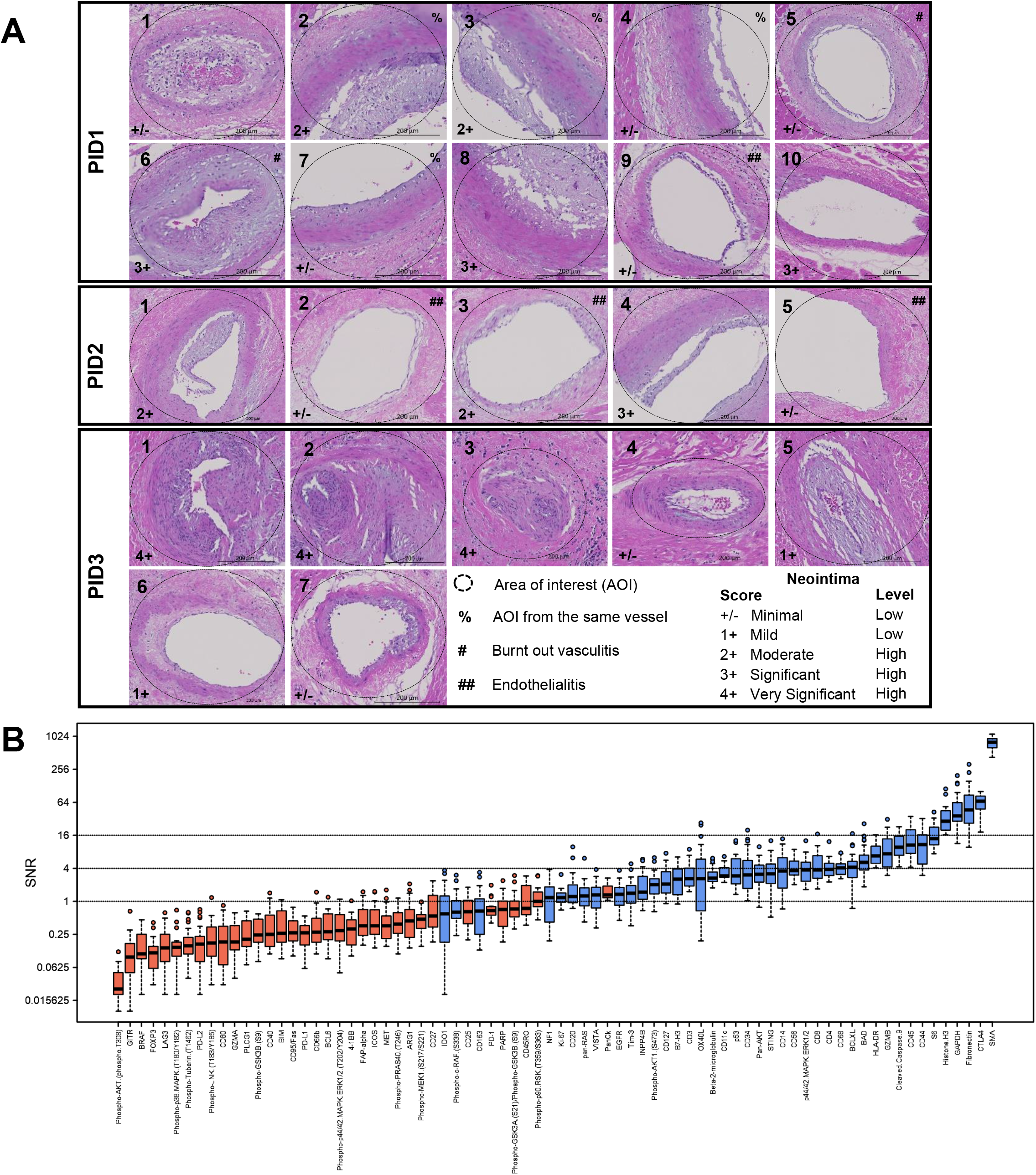
Vessel regions from cardiac allograft vasculopathy (CAV) rejected grafts exhibit varying degrees of neointimal thickening and express 41 protein markers. **A)** A total of 22 arterial areas of interest (AOIs) from CAV+DSA+ patients (PID1-3; 2 females and 1 male) were subjected to protein and RNA digital spatial profiling (DSP). In total, 11 AOIs were scored with ‘low’ neointima (+/- minimal and 1+ mild) and 11 with ‘high’ neointima (2+ moderate, 3+ significant, and 4+ very significant) by H&E staining (bar=200µm). **B)** Protein profiling identified a total of 41 protein markers which were similarly expressed across all AOIs (average SNR>1 in blue). Protein data was normalized to the average of negative IgG controls and reported as a signal to noise ratio (SNR). Proteins with an average SNR<1 (in red) were considered not expressed.

### 5.2. Top transcripts in arteries encode for DSA-mediated immune responses and vascular remodeling

Transcriptomic analysis identified a total of 10,746 genes with an average SNR>1 and coefficient of variation (COV)>0.19 across all 22 arterial AOIs. A total of 74 transcripts had an average SNR>10 (excluding ribosomal and ATP genes) (**Figure 2A**). Mainly, *ADAM15* (a disintegrin and metalloprotease) which has been shown to mediate endothelial hyperpermeability during inflammation was highly expressed (median SNR 96.5)^18^. Immunoglobulin transcripts (e.g., *IGKC* and *IGHG1/2/3/4*) were also highly elevated. This was accompanied by the expression of HLA class I (*HLA-B*), IFN-у inducible HLA class II molecules (e.g., *HLA-DRB1* and *HLA-DRA*), and HLA class II chaperone transcripts (*CD74*)^19^. AOIs also exhibited a relatively high expression of transcripts described in vascular remodeling, angiogenesis, platelet activation, cell proliferation, migration, and immune infiltration (*IGFBP7*, *TMSB4X*, *FLNA*, *TIMP1*, *MAZ*, *CRYAB*, *TMEM106C*) ^20–25^. Metascape gene enrichment analysis of the top 231 transcripts (average SNR>5) revealed that AOIs were enriched in pathways relating to EC angiogenesis (‘*VEGFA-VEGFR2 signaling pathway’*), DSA activation (‘*immunoglobulin mediated immune responses’*), thrombosis (‘*platelet degranulation’*), and pro-inflammatory cytokines (‘*IL-18 signaling pathway’*) (**Figure 2B**). Highly expressed transcripts most likely were attributed to fibroblast, ECs, macrophages, and memory CD8 T-cells as these were most prominent cell types by cellular abundance counts (**Supplementary Figure 2A**). Cell abundance counts upheld protein results as CD68 and CD8 were among the top immune-cell markers. Specifically, CD8 protein expression significantly correlated with CD8 memory abundance counts (and not CD8 naïve T-cells) further suggesting the presence of memory T-cells (**Supplementary Figure 2B**). Additionally, we identified that CD45, CD8, CD56, CD20, CD127, and CD11c protein markers significantly correlated with RNA counterpart expression validating the level of expression of these cell types in each AOI (**Figure 2C**).

**Figure 2.**
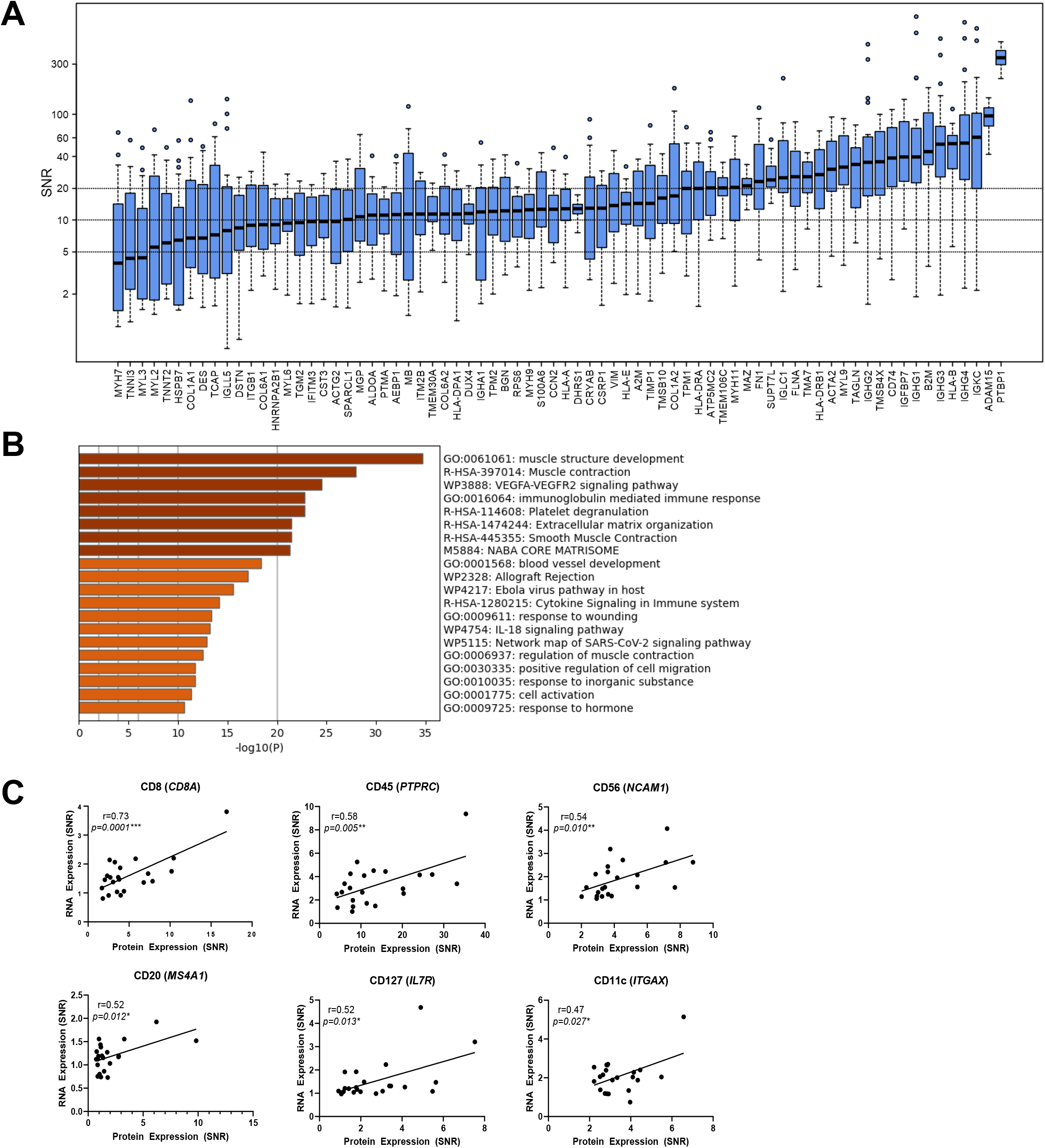
Top transcripts in arterial AOIs encode for DSA-mediated immune responses and vascular remodeling. **A)** Expression levels of the top 74 transcripts (average SNR>10) ranked by median values (excluding ribosomal and ATP transcripts). Whole genome RNA data was normalized to the average (+2SD) of internal negative controls (NegProbe-WTX) and is reported as a signal to noise ratio (SNR). **B)** Pathway enrichment analysis of the top 231 transcripts (with average SNR>5) using Metascape. **C)** The protein expression of CD8, CD45, CD56, CD20, CD127, and CD11c exhibited a significantly positive correlation with RNA counterpart expression. Statistical significance was determined using Pearson correlation test (*p<0.05* (*), *p<0.01*(**), *p<0.001*(***).

### 5.3. Arterial AOIs with low neointima exhibit higher inflammation and cell death while AOIs with high neointima exhibit remodeling and fibrotic profiles

Although all arteries were found to exhibit similar protein markers of both innate and adaptive immune infiltrates, we questioned whether the degree of inflammation varied between AOIs scored with a low (+/- minimal and 1+ mild) or high (2+ moderate, 3+ significant, and 4+ very significant) neointima (N=11 AOIs for each condition). Unsupervised clustering of the 41 proteins (SNR>1) illustrated that AOIs with low neointima clustered closer together and displayed higher protein expression (**Figure 3A**). This was also observed when clustering AOIs for each individual patient (**Supplementary Figure 3**). A total of 8 differentially expressed proteins (DEPs) were increased in arteries with low neointima (**Figure 3B**). These included markers involved in T-cell clonal expansion/survival (TNFSF4/OX40L), memory T-cells (CD45RO and CD127), checkpoint inhibitors (TIM-3), monocytes (CD14), regulators of PI3K/AKT signaling (INPP4B), MAPK signaling (pan-Ras), and pro-apoptotic proteins (p53) (**Table 1**). AOIs with low neointima also upregulated *VWF* (indicative of EC activation)^26^. Meanwhile, AOIs with high neointima significantly increased TGFβ-regulated genes (*CSRP1* and *TAGLN)* and adipocyte differentiation factor (*ADIRF/APM2*) (**Figure 3C and Table 1**)^27–29^.

**Figure 3.**
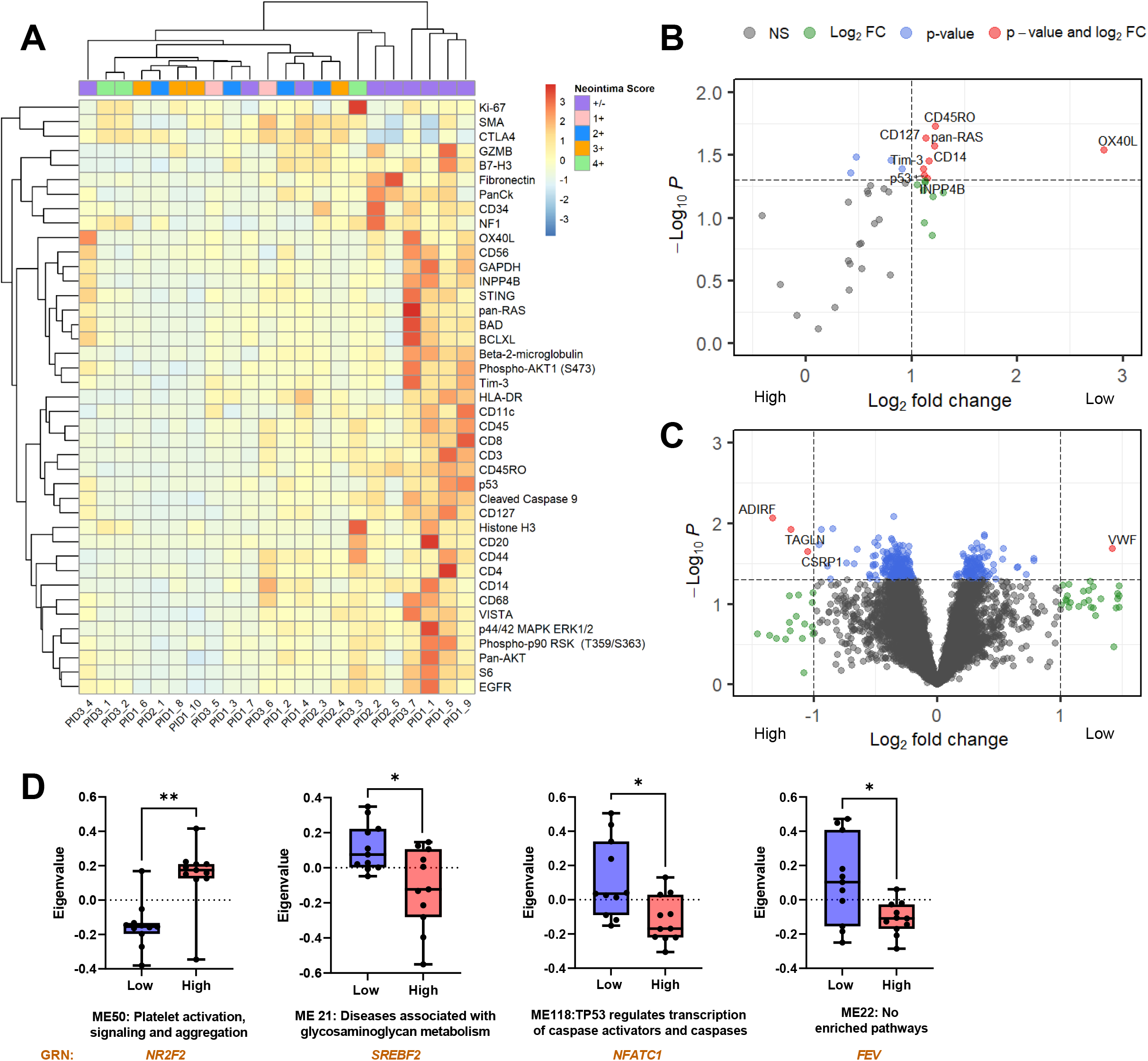
Arterial AOIs containing high and low neointima exhibit differences in protein and RNA expression. **A)** Unsupervised clustering of the 41 expressed proteins (average SNR>1) across all 22 arterial AOIs. **B)** Differentially expressed proteins (DEPs) (in red dots) between AOIs with low and high neointima. **C)** Differentially expressed genes (DEGs) between AOIs with low and high neointima (in red dots). **D)** Weighted gene co-expression network analysis (WGCNA) was used to generate gene modules (ME) of co-expressed genes. Expression of each module is represented by eigengene values (Eigenvalue). Module names were defined by Reactome pathway enrichment analysis. Arterial AOIs with low neointima increased ME21, ME22, and ME118 while AOIs with high neointima increased ME50. The highest positively correlated gene regulatory network (GNR) for each module is included below (orange text). Statistical significance for volcano plots (Log_2_FC>1 and *p-value*<*0.05*) and for comparing modules (Eigenvalues) was determined using linear mixed-effects model (including PID as random effects) using in R software (*p<0.05* (*) and *p<0.01* (**)).

To further explore transcriptional differences between low and high neointima AOIs, we utilized whole gene co-expression network analysis (WGCNA) to define modules of co-expressed transcripts^15^. AOIs with low neointima upregulated three modules (ME21, ME22, and ME118), two of which were enriched for ‘*Activation of matrix metalloproteinases*’ and ‘*TP53 regulation of caspase activators and caspases’*. High neointima AOIs upregulated one module (ME50) enriched for pathways involved in ‘*Platelet activation, signaling and aggregation*’ (**Figure 3D and Supplementary Table 6**). Gene regulatory network (GRN) analysis enabled the identification of key transcription factors correlating co-expression modules. Specifically, ME21 and ME118 highly correlated with inflammatory driven transcription factors *SREBF2* and *NFATC1*, respectively^30, 31^. Meanwhile, ME50 highly correlated with *NR2F2*, which promotes cell proliferation and TGF-β-dependent epithelial-to-mesenchymal transition (EMT) (**Figure 3D and Supplementary Table 6**)^32^.

In addition, we identified that numerous protein markers encoding inflammatory infiltrates (e.g., CD45, CD44, CD8, OX40L, GZMB) and apoptosis (p53 and BAD) highly correlated with co-expressing modules enriched for ‘*interferon gamma signaling*’, ‘*Toll-like receptor cascades*’, ‘*NF-kB phosphorylation*’ and ‘*TP53 regulation of caspase activators and caspases*’ (ME65, 77, 91, 118, 242, and 304) only in AOIs with low neointima. Alternatively, proteins associated with growth factor receptors (EGFR), regulators of cell growth/division (NF1), smooth muscle cell markers (SMA), and cell survival proteins (BCL-XL) significantly correlated with modules enriched for ‘*MAPK1/MAPK3 signaling*’, ‘*signaling by FGFR*’, ‘*FGFR2 ligand binding and activation*’, and ‘*VEGFA-VEGFR2 pathway*’ (ME161, 205, 206) only in AOIs with high neointima (**Table 2 and supplementary Figure 4**). Cellular deconvolution reinforced neointimal differences since AOIs with high neointima exhibited a significantly higher number of fibroblasts, while AOIs with low neointima contained a higher number of classical monocytes (**Supplementary Figure 2C**).

### 5.4. CAV lesion progression is associated with distinct gene expression signatures and gene regulatory networks

To identify genes associated with different stages of neointima, we performed a linear regression of neointima score on normalized expression of each gene. From principal component analysis of genes significantly associated with neointima score (n=245), we identified the first principal component (PC1) as increasing in a stepwise manner with increasing neointima score (**Figure 4A**). Finally, to identify key genes and GRNs which may be driving vessel occlusion, we tested for genes/GRNs associated with PC1 score. A total of 36 genes were significantly associated with PC1 by linear regression analysis (adjusted *p-value* < 0.05) (**Figure 4B and Supplementary Table 7**). Specifically, arteries with mild neointima (+/-) contained heat shock proteins (*CRYAB/HSBP5* and *HSPB7*) which increase in response to inflammatory stress^25, 33^. Vessels with minimal neointima (1+) increased the antigen T-cell receptor (*TARP*) which promotes tumor cell proliferation and migration^34^. Vessels with moderate neointima (2+) increased a set of genes described in remodeling and EMT (*TAGLN*, *CSRP1, and FLNA*)^22, 27, 28^. Vessels with significant neointima (3+) only increased ubiquitin and RNA-binding protein (*UBAP2L*), a principal mediator of EMT/fibrosis and emerging biomarker and therapeutic target of EndoMT in cancer^35, 36^. Finally, arteries with very significant neointima (4+) increased early growth response-1 (*EGR1*) and *FOS* transcription factors, implicated in cancer progression and fibrosis^37, 38^. Notably, transcripts involved in inhibiting tumor cell proliferation, EMT/fibrosis (*OGN*) and anti-inflammatory regulators (*ZFP36*) were also identified in vessels with moderate (2+) and very significant neointima (4+), respectively^39, 40^. Major GRNs associated with increasing neointima also included transcription factors involved promoting cell proliferation, migration, EMT/EndoMT, fibrosis (*NF2F2, JUN, HEY, ATF3, ATF4, and FOXP1)* and angiogenesis (*RUNX1*)^32, 41–46^ (**Figure 4C**).

**Figure 4.**
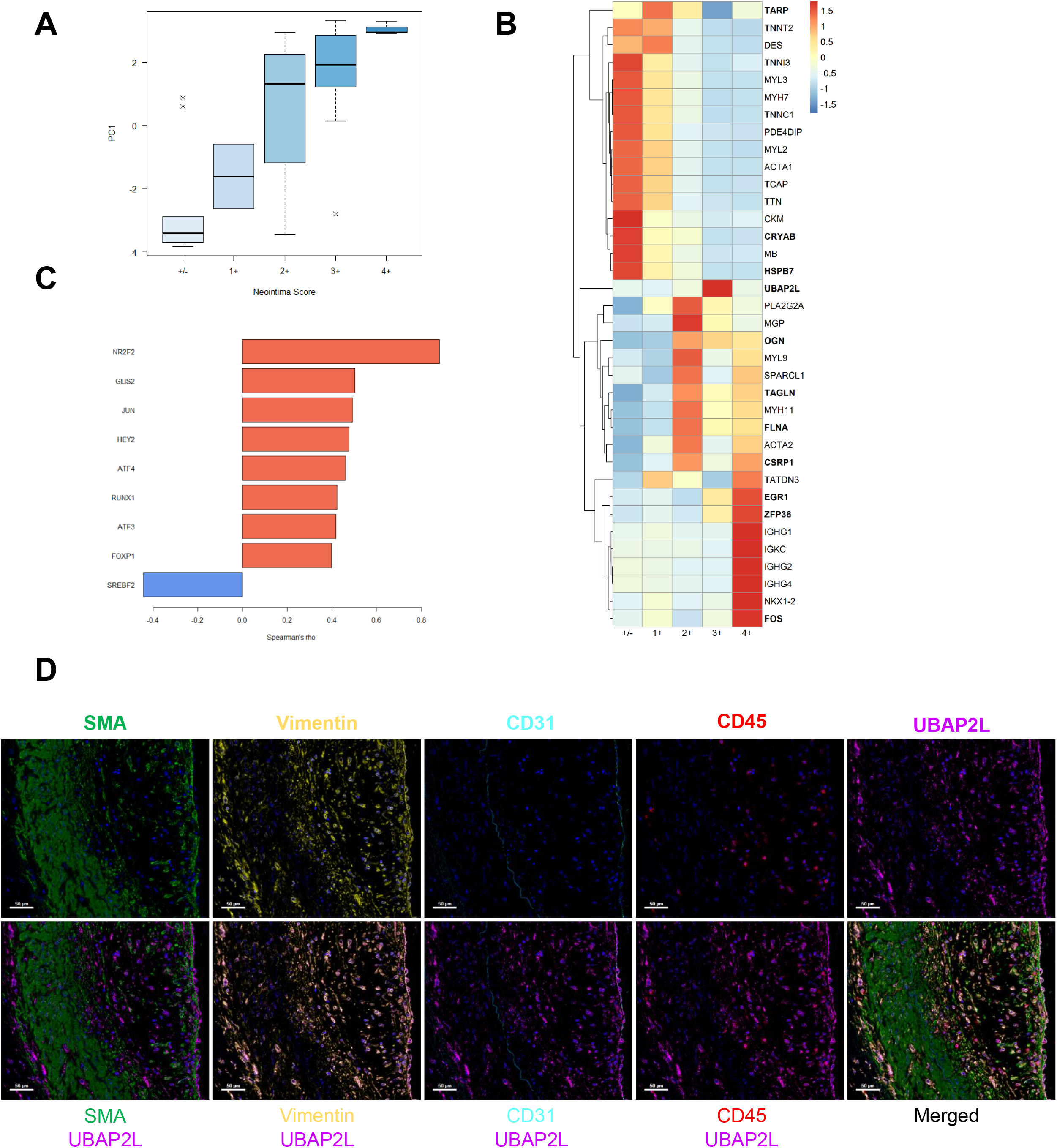
A total of 36 genes and 9 gene regulatory networks (GNRs) were significantly associated with neointima score. **A)** From principal component analysis of genes significantly associated with neointima score (n=245), we identified the first principal component (PC1) as increasing in a stepwise manner with increasing neointima score. **B)** A total of 36 genes were significantly associated (Adjusted *p-value* < *0.05*) with PC1 showed distinct patterns with increasing neointima score. Genes in bold have been characterized to promote cell proliferation, migration, epithelial-to-mesenchymal transition (EMT) and fibrosis (p-values found in **Supplementary Table 7**). **C)** A total of 9 gene regulatory networks (GNRs) correlated with PC1 and therefore increasing neointima score (*p-value* < *0.05*). **D)** UBAP2L co-localized with CD31+ ECs and Vimentin+ mesenchymal cells within the neointima. Representative image shows PID1 AOI 8 which exhibits significant neointima (3+) (scale bar=50µm).

Neointima driven genes resided within scores of +/-, 2+, and 4+, suggesting sequential stages in which genes may be turned on-and-off. Notably, UBAP2L was only increased in AOIs with a score of 3+. Since UBAP2L has been reported as a key mediator of EMT in cancer, we questioned whether it was also expressed in arterial AOIs as a possible mediator of EndoMT. Multiplex immunofluorescent staining of cardiac explants revealed that UBAP2L co-localized with CD31+ ECs and Vimentin+ mesenchymal cells. CD31+ ECs also co-expressed Vimentin suggesting activation toward mesenchymal phenotype. UBAP2L did not co-localize with SMA+ or CD45+ cells (**Figure 4D**). A total of 16/22 (72%) of arterial AOIs expressed UBAP2L co-localized within CD31+Vimentin+ ECs and CD31-Vimentin+ mesenchymal cells. Specifically, 9/11 (81%) high neointima AOIs while only 7/11 (63%) of low neointima AOIs expressed UBAP2L.

## 6. Discussion

In this study, we utilized GeoMx DSP technology to examine both the protein and transcriptomic expression of arterial AOIs from HLA DSA+CAV+ rejected cardiac explants. Among the 22 arterial regions examined, we identified similarly expressed protein markers relating to innate and adaptive immune cells, cell activation, and cell death. Whole transcriptome analysis highlighted transcripts involved in DSA-mediated responses and vascular remodeling. Moreover, we demonstrated that AOIs with low neointima exhibit higher inflammatory and cell death profiles while AOIs with high neointima exhibit lesions undergoing proliferation, migration, and EndoMT/fibrosis. Finally, we identified key genes (e.g., *UBAP2L*) and transcription factors associated with increasing neointima scores, many implicated in promoting proliferation, migration, and EMT/fibrosis. By examining the similarities and heterogeneities between arterial regions, we present a unique approach to further delineate the potential stages of CAV progression.

Macrophages and T cells have been found to accumulate within the neointima of human CAV affected vessels^47, 48^. This is concordant with our study as CD68+ and CD4/CD8+ markers were the top immune infiltrates in both protein and RNA cell abundance datasets. Monocyte recruitment may occur in the early phases of CAV as we observed significantly higher protein expression of CD14 and classical monocytes counts in arteries with low neointima. HLA DSA can rapidly increase intracellular calcium and endothelial presentation of P-selectin, which supports monocyte binding^7, 49^. Notably, CD68 protein expression also significantly correlated with co-expressing module ME206 in AOIs with high neointima. ME206 was enriched for pathways related to ‘*Fc gamma receptor dependent phagocytosis*’ indicative of polarized macrophage specific functions within developing lesions^50^.

Endothelial cell injury is a major hallmark of AMR and in the early phases of CAV. Specifically, high protein expression of CD8+ T cells, CD44 activation marker, cleaved caspase 9, and GZMB together suggest cell-targeted injury via the perforin/granzyme apoptosis pathway by cytotoxic T-cells. Studies using minor MHC mismatched heterotopic cardiac transplants showed that GZMB knockout recipient mice exhibited a significant reduction in luminal narrowing of allograft arteries compared to wildtype recipients^51^. Human coronary arteries with advanced atherosclerosis and CAV lesions have also been shown to exhibit high levels of GZMB localized within infiltrating leukocytes underlying the endothelium, in the deep intima, and in perivascular infiltrates in the adventitia^52^. GZMB-induced cell apoptosis can be enhanced by the pro-apoptotic protein BAD (BCL-2 associated agonist of cell death) or hindered by the anti-apoptotic BCL-2 family protein, BCL-XL. These regulatory proteins shared similar protein expression levels to GZMB and exhibited the highest significant correlation with each other. This is likely because activated BAD binds to and inhibits the anti-apoptotic function of BCL-XL^53^ Expression of EC anti-apoptotic proteins can be potentiated by antibody ligation of HLA class I and II molecules which activate the PI3K/AKT pathway and upregulation of BCL-2 and BCL-XL^54^. Hence, ECs may acquire anti-apoptotic defense mechanisms that promote EndoMT via HLA outside in signaling.

Arterial AOIs also showed high expression of transcripts associated with DSA-mediated responses including multiple immunoglobulins (IgGs), suggesting the presence of antibody secreting cells such as B cells and/or plasma cells. Since CD20 protein expression across AOIs was relatively low (SNR=1.21), it is likely that these IgG transcripts are encoded by mature plasma cells (which decrease CD20 expression). This was confirmed in a study by Chattejee et al., in which majority of the CD138+ differentiated plasma cells around CAV affected coronary arteries from rejected explants secreted IgG^55^. Moreover, microarray studies by Wehner et al., have shown a strong immunoglobulin transcriptome in CAV rejected explants in contrast to atherosclerosis^56^.

All arteries shared high expression for transcripts associated with EC and SMC activation, remodeling, angiogenesis, and platelet activation^20–24^. We compared transcripts with an average SNR > 5 to those identified in pathological AMR+ DSA+ EMBs reported by Loupy et al. using microarray technology^11^. Similarly, we detected the expression of IFN-y inducible genes (HLA class II transcripts), and NK-cell activation (*FCGR3A*). However, our results may not fully encompass the transcriptomic profile described by studies using EMBs due to anatomical differences within the heart. These differences were emphasized in a mouse model of chronic AMR in which the recipient heart apical tissue had 51 significant upregulated genes (including activated immune infiltrates) compared to two upregulated genes (E-selectin and TLR4) in epicardial CAV lesion^57^.

Multi-omics analysis between AOIs with low and high neointima proposes a sequential time-frame of the vascular changes leading to CAV (**Supplementary Figure 5**). Specifically, ‘early’ lesions exhibit higher inflammatory profiles consisting of lymphocyte and monocyte infiltrates, while ’late’ lesions contain higher proliferative and fibromuscular/mesenchymal tissue with lower inflammation. These observations are in accordance with the results of Huibers et al. who identified three histopathological patterns of CAV characterized by inflammatory lesions, increased SMCs, followed by fibrotic lesions^58^. AOIs with low neointima mainly exhibit features of burnt-out vasculitis (thickening of the vessel wall) or endothelialitis (cells beneath the endothelium). Based on the significantly higher memory T-cell and cell death markers in low neointima AOIs, we speculate that T-cells infiltrate subendothelial regions where they proliferate and mediate EC-injury. The accumulation of T-cells expressing perforin-containing granules in the subendothelial region of early CAV lesions has been previously reported^59^. OX40L (*TNFRSF4*) was the highest DEP between low and high neointima AOIs. OX40L (expressed on APCs, NK cell, activated CD4 T-cells, ECs and mast cells) binds to OX40+ antigen activated T-cells stimulating T-cell proliferation, clonal expansion, survival, and effector cytokine release^60^. Studies using mouse heart transplant models have highlighted the benefits of OX40L blockade in promoting graft survival by inducing Tregs^61^. The notion that AOIs with low neointima exhibit characteristics of early vascular inflammation and cell death was furthered reinforced by a significant increase in *VWF*, and in ME118 (enriched for TP53 activation of caspases). Moreover, ME118 significantly correlated with nuclear factor of activated T-cells (*NFATC1*), a main driver of vascular inflammation, and regulator of cytotoxic CD8+ T-cells^31^.

Alternatively, AOIs with high neointima mainly demonstrated features of ongoing neointima expansion as seen by an increase in transcripts involved in myofibroblast differentiation (*CSRP1*), and remodeling (*TAGLN*)^27, 28^. AOIs with high neointima upregulated pathways related to ‘*Platelet activation, signaling and aggregation*’ (ME50). DSA crosslinking to MHC I antigens induces Weibel-Palade bodies (WPb) exocytosis of P-selectin and vWF which can increase platelet and leukocyte infiltration which aggravate vessel pathology^7, 49, 62, 63^.

High neointima AOIs also increased GRNs involved in cell proliferation (*JUN*), EndoMT (*NR2F2*, *HEY2*, and *FOXP1*), angiogenesis (*RUNX1*) and fibrosis (*ATF3* and *ATF4*)^27, 32, 41–46^. Notably, DEGs (*CSRP1* and *TAGLN*) and GRNs (*NR2F2*, *ATF3*, *FOXP1*) encompassed genes which promote EndoMT by regulating or being induced by TGF-β^27, 28, 32, 43^. Similar findings were observed by linear regression analysis as key genes involved tumor cell proliferation, migration, EMT/fibrosis, and anti-inflammatory modulators (*ZFP36*) highly associated with increasing neointima scores. Intriguingly, inhibitors of EMT/fibrosis were also identified in high neointima AOIs in both GRN (*GLIS2*) and linear regression (*OGN*) results^39, 64^. This raises potential targetable mechanisms which may help delay CAV progression.

Furthermore, we identified UBAP2L as a new potential regulator of EndoMT in CAV. UBAP2L is a major regulator of EMT via numerous mechanisms. This includes positively regulating the transcription factor SNAIL1 which suppresses E-cadherin expression and by sustaining cell proliferation via regulating cyclins and PI3K/Akt signaling^35, 36^. Cancer cells overexpressing UBAP2L are characterized by upregulating mesenchymal factors (N-cadherin and Vimentin). Our findings suggest that CD31+Vimentin+UBAP2L+ ECs may be in an intermediate stage of EndoMT. AOIs with high neointima also contained CD31-Vimentin+UBAP2L+ mesenchymal cells which may be at a more terminal stage of EndoMT but still undergoing proliferation and growth possibly induced by UBAP2L.

Finally, neointimal differences became more apparent as the majority of markers encoding effector immune cell infiltrates significantly correlated with inflammatory co-modules enriched for IFN-y signaling, TLR cascades, and NF-kB phosphorylation. Meanwhile, the majority of protein markers associated with cell growth factors and proliferation (e.g., EGFR and NF1) correlated with profibrotic modules enriched for FGFR, FDFR2, and VEGFA-VEGFR2 signaling (**Table 3**). Antibody ligation of HLA I and HLA II molecules stimulates EC cell proliferation and migration via activation of PI3K/AKT, ERK, and mTOR signaling^4, 5, 65^. Moreover, anti-HLA I antibodies can mediate an increase in FGFR cell surface expression^66^.Together, these findings reinforce the notion that ECs and SMCs undergo active proliferative, migrative, and pro-fibrotic signals ultimately contributing to vessel occlusion.

In summary, our data suggest a temporal progression whereby initial inflammation by mononuclear cell infiltrates in the intima later promote proliferative and fibrotic responses (**Figure 5**). Although it is difficult to predict whether AOIs with low neointima will indeed progress into arteries with high neointima, our findings emphasize on the degree of arterial heterogeneity in CAV rejected explants. Similar observations were reported in a pathological study by Lu et al., in which vessels from 64 CAV rejected explants exhibited diverse characteristics such as intimal fibromuscular hyperplasia, vasculitis, and endothelialitis^48^. Prior studies have also pointed to a possible relationship between AMR and EMBs exhibiting vascular endothelialitis in developing CAV^67^.

While our study reveals novel protein and transcriptomic profiles in arteries from DSA+CAV+ rejected cardiac explants, there remains a few limitations to be addressed. First, DSP analysis represents data from a selected region rather than at the single-cell level. On-going studies are focused on identifying cell-type specific proteins and transcripts. We encountered variability between patient characteristics yet found no significant correlations between patient age at transplant and neointima score nor with time-post transplant and neointima score. Third, additional studies characterizing patient explants from DSA negative controls are needed. Future studies will focus on validating the expression of highly expressed transcripts in comparison with DSA negative controls. Finally, our results feature novel differences between low and high neointima AOIs however, further longitudinal studies would be needed to confirm the fate of low neointimal arteries.

To our knowledge, this is the first spatial multi-omics study examining arterial vessels with varying degree of neointima formation in DSA+CAV+ cardiac transplant rejected explants. Our findings further accentuate on the degree of vessel heterogeneity and profiles not usually identified by pathology alone. We anticipate this newly generated dataset can serve as resource for future high throughput or basic studies investigating the mechanisms of DSA mediated injury and CAV development.

## 7. Acknowledgements

We would like to thank the NanoString Technology Access Program (TAP) for this opportunity and UCLA TPCL for their resources. Finally, the authors thank the organ donors and their families for their generous gifts of life and knowledge.

## 9. Author Contributions

JNM and EFR designed research study. JNM and HCP performed data analysis. RAS, GAF, and WMB identified patient tissue, region selection, and grading. WMB and RLF contributed to the design of the study. JNM and EFR wrote the manuscript. All authors read and approved the manuscript.

## Non-standard Abbreviations and Acronyms

CAV: cardiac allograft vasculopathy
AMR: antibody-mediated Rejection
DSA: donor specific HLA antibodies
EC: Endothelial cell
HLA: human leukocyte antigen
AOIs: areas of interest
EMT: Epithelial-to-mesenchymal transition
EndoMT: Endothelial-to-mesenchymal transition

## 11. References

1. Khush KK, Cherikh WS, Chambers DC, Goldfarb S, Hayes D, Jr., Kucheryavaya AY, Levvey BJ, Meiser B, Rossano JW, Stehlik J. The International Thoracic Organ Transplant Registry of the International Society for Heart and Lung Transplantation: Thirty-fifth Adult Heart Transplantation Report-2018; Focus Theme: Multiorgan Transplantation. J Heart Lung Transplant. 2018;37:1155–1168. doi: 10.1016/j.healun.2018.07.022

2. Jansen MA, Otten HG, de Weger RA, Huibers MM. Immunological and Fibrotic Mechanisms in Cardiac Allograft Vasculopathy. Transplantation. 2015;99:2467–2475. doi: 10.1097/tp.0000000000000848

3. Colvin MM, Cook JL, Chang P, Francis G, Hsu DT, Kiernan MS, Kobashigawa JA, Lindenfeld J, Masri SC, Miller D, et al. Antibody-mediated rejection in cardiac transplantation: emerging knowledge in diagnosis and management: a scientific statement from the American Heart Association. Circulation. 2015;131:1608–1639. doi: 10.1161/cir.0000000000000093

4. Zhang X, Rozengurt E, Reed EF. HLA class I molecules partner with integrin β4 to stimulate endothelial cell proliferation and migration. Sci Signal. 2010;3:ra85. doi: 10.1126/scisignal.2001158

5. Jin YP, Valenzuela NM, Zhang X, Rozengurt E, Reed EF. HLA Class II-Triggered Signaling Cascades Cause Endothelial Cell Proliferation and Migration: Relevance to Antibody-Mediated Transplant Rejection. J Immunol. 2018;200:2372–2390. doi: 10.4049/jimmunol.1701259

6. Salehi S, Sosa RA, Jin YP, Kageyama S, Fishbein MC, Rozengurt E, Kupiec-Weglinski JW, Reed EF. Outside-in HLA class I signaling regulates ICAM-1 clustering and endothelial cell-monocyte interactions via mTOR in transplant antibody-mediated rejection. Am J Transplant. 2018;18:1096–1109. doi: 10.1111/ajt.14544

7. Jin YP, Nevarez-Mejia J, Terry AQ, Sosa RA, Heidt S, Valenzuela NM, Rozengurt E, Reed EF. Cross-Talk between HLA Class I and TLR4 Mediates P-Selectin Surface Expression and Monocyte Capture to Human Endothelial Cells. J Immunol. 2022;209:1359–1369. doi: 10.4049/jimmunol.2200284

8. Thomas KA, Valenzuela NM, Reed EF. The perfect storm: HLA antibodies, complement, FcγRs, and endothelium in transplant rejection. Trends Mol Med. 2015;21:319–329. doi: 10.1016/j.molmed.2015.02.004

9. Tellides G, Pober JS. Interferon-gamma axis in graft arteriosclerosis. Circ Res. 2007;100:622–632. doi: 10.1161/01.Res.0000258861.72279.29

10. Lu X, Gong J, Dennery PA, Yao H. Endothelial-to-mesenchymal transition: Pathogenesis and therapeutic targets for chronic pulmonary and vascular diseases. Biochem Pharmacol. 2019;168:100–107. doi: 10.1016/j.bcp.2019.06.021

11. Loupy A, Duong Van Huyen JP, Hidalgo L, Reeve J, Racapé M, Aubert O, Venner JM, Falmuski K, Bories MC, Beuscart T, et al. Gene Expression Profiling for the Identification and Classification of Antibody-Mediated Heart Rejection. Circulation. 2017;135:917–935. doi: 10.1161/circulationaha.116.022907

12. Halloran PF, Potena L, Van Huyen JD, Bruneval P, Leone O, Kim DH, Jouven X, Reeve J, Loupy A. Building a tissue-based molecular diagnostic system in heart transplant rejection: The heart Molecular Microscope Diagnostic (MMDx) System. J Heart Lung Transplant. 2017;36:1192–1200. doi: 10.1016/j.healun.2017.05.029

13. Mantell BS, Cordero H, See SB, Clerkin KJ, Vasilescu R, Marboe CC, Naka Y, Restaino S, Colombo PC, Addonizio LJ, et al. Transcriptomic heterogeneity of antibody mediated rejection after heart transplant with or without donor specific antibodies. J Heart Lung Transplant. 2021;40:1472–1480. doi: 10.1016/j.healun.2021.06.012

14. Merritt CR, Ong GT, Church SE, Barker K, Danaher P, Geiss G, Hoang M, Jung J, Liang Y, McKay-Fleisch J, et al. Multiplex digital spatial profiling of proteins and RNA in fixed tissue. Nat Biotechnol. 2020;38:586–599. doi: 10.1038/s41587-020-0472-9

15. Langfelder P, Horvath S. WGCNA: an R package for weighted correlation network analysis. BMC Bioinformatics. 2008;9:559. doi: 10.1186/1471-2105-9-559

16. Aibar S, González-Blas CB, Moerman T, Huynh-Thu VA, Imrichova H, Hulselmans G, Rambow F, Marine JC, Geurts P, Aerts J, et al. SCENIC: single-cell regulatory network inference and clustering. Nat Methods. 2017;14:1083–1086. doi: 10.1038/nmeth.4463

17. Danaher P, Kim Y, Nelson B, Griswold M, Yang Z, Piazza E, Beechem JM. Advances in mixed cell deconvolution enable quantification of cell types in spatial transcriptomic data. Nat Commun. 2022;13:385. doi: 10.1038/s41467-022-28020-5

18. Sun C, Wu MH, Guo M, Day ML, Lee ES, Yuan SY. ADAM15 regulates endothelial permeability and neutrophil migration via Src/ERK1/2 signalling. Cardiovasc Res. 2010;87:348–355. doi: 10.1093/cvr/cvq060

19. Su H, Na N, Zhang X, Zhao Y. The biological function and significance of CD74 in immune diseases. Inflamm Res. 2017;66:209–216. doi: 10.1007/s00011-016-0995-1

20. Lisowska A, Święcki P, Knapp M, Gil M, Musiał WJ, Kamiński K, Hirnle T, Tycińska A. Insulin-like growth factor-binding protein 7 (IGFBP 7) as a new biomarker in coronary heart disease. Adv Med Sci. 2019;64:195–201. doi: 10.1016/j.advms.2018.08.017

21. Rossdeutsch A, Smart N, Dubé KN, Turner M, Riley PR. Essential role for thymosin β4 in regulating vascular smooth muscle cell development and vessel wall stability. Circ Res. 2012;111:e89–102. doi: 10.1161/circresaha.111.259846

22. Bandaru S, Ala C, Zhou AX, Akyürek LM. Filamin A Regulates Cardiovascular Remodeling. Int J Mol Sci. 2021;22. doi: 10.3390/ijms22126555

23. Barton PJ, Birks EJ, Felkin LE, Cullen ME, Koban MU, Yacoub MH. Increased expression of extracellular matrix regulators TIMP1 and MMP1 in deteriorating heart failure. J Heart Lung Transplant. 2003;22:738–744. doi: 10.1016/s1053-2498(02)00557-0

24. Smits M, Wurdinger T, van het Hof B, Drexhage JA, Geerts D, Wesseling P, Noske DP, Vandertop WP, de Vries HE, Reijerkerk A. Myc-associated zinc finger protein (MAZ) is regulated by miR-125b and mediates VEGF-induced angiogenesis in glioblastoma. Faseb j. 2012;26:2639–2647. doi: 10.1096/fj.11-202820

25. Zhang J, Liu J, Wu J, Li W, Chen Z, Yang L. Progression of the role of CRYAB in signaling pathways and cancers. Onco Targets Ther. 2019;12:4129–4139. doi: 10.2147/ott.S201799

26. Ruggeri ZM. The role of von Willebrand factor in thrombus formation. Thromb Res. 2007;120 Suppl 1:S5–9. doi: 10.1016/j.thromres.2007.03.011

27. Järvinen PM, Myllärniemi M, Liu H, Moore HM, Leppäranta O, Salmenkivi K, Koli K, Latonen L, Band AM, Laiho M. Cysteine-rich protein 1 is regulated by transforming growth factor-β1 and expressed in lung fibrosis. J Cell Physiol. 2012;227:2605–2612. doi: 10.1002/jcp.23000

28. Elsafadi M, Manikandan M, Dawud RA, Alajez NM, Hamam R, Alfayez M, Kassem M, Aldahmash A, Mahmood A. Transgelin is a TGFβ-inducible gene that regulates osteoblastic and adipogenic differentiation of human skeletal stem cells through actin cytoskeleston organization. Cell Death Dis. 2016;7:e2321. doi: 10.1038/cddis.2016.196

29. Scott BJ, Qutob S, Liu QY, Ng CE. APM2 is a novel mediator of cisplatin resistance in a variety of cancer cell types regardless of p53 or MMR status. Int J Cancer. 2009;125:1193–1204. doi: 10.1002/ijc.24465

30. Dong G, Huang X, Wu L, Jiang S, Tan Q, Chen S. SREBF2 triggers endoplasmic reticulum stress and Bax dysregulation to promote lipopolysaccharide-induced endothelial cell injury. Cell Biol Toxicol. 2022;38:185–201. doi: 10.1007/s10565-021-09593-1

31. Klein-Hessling S, Muhammad K, Klein M, Pusch T, Rudolf R, Flöter J, Qureischi M, Beilhack A, Vaeth M, Kummerow C, et al. NFATc1 controls the cytotoxicity of CD8(+) T cells. Nat Commun. 2017;8:511. doi: 10.1038/s41467-017-00612-6

32. Wang H, Nie L, Wu L, Liu Q, Guo X. NR2F2 inhibits Smad7 expression and promotes TGF-β-dependent epithelial-mesenchymal transition of CRC via transactivation of miR-21. Biochem Biophys Res Commun. 2017;485:181–188. doi: 10.1016/j.bbrc.2017.02.049

33. Finka A, Mattoo RU, Goloubinoff P. Experimental Milestones in the Discovery of Molecular Chaperones as Polypeptide Unfolding Enzymes. Annu Rev Biochem. 2016;85:715–742. doi: 10.1146/annurev-biochem-060815-014124

34. Vanhooren J, Derpoorter C, Depreter B, Deneweth L, Philippé J, De Moerloose B, Lammens T. TARP as antigen in cancer immunotherapy. Cancer Immunol Immunother. 2021;70:3061–3068. doi: 10.1007/s00262-021-02972-x

35. Ye T, Xu J, Du L, Mo W, Liang Y, Xia J. Downregulation of UBAP2L Inhibits the Epithelial-Mesenchymal Transition via SNAIL1 Regulation in Hepatocellular Carcinoma Cells. Cell Physiol Biochem. 2017;41:1584–1595. doi: 10.1159/000470824

36. Li Q, Wang W, Hu YC, Yin TT, He J. Knockdown of Ubiquitin Associated Protein 2-Like (UBAP2L) Inhibits Growth and Metastasis of Hepatocellular Carcinoma. Med Sci Monit. 2018;24:7109–7118. doi: 10.12659/msm.912861

37. Bhattacharyya S, Wu M, Fang F, Tourtellotte W, Feghali-Bostwick C, Varga J. Early growth response transcription factors: key mediators of fibrosis and novel targets for anti-fibrotic therapy. Matrix Biol. 2011;30:235–242. doi: 10.1016/j.matbio.2011.03.005

38. Milde-Langosch K. The Fos family of transcription factors and their role in tumourigenesis. Eur J Cancer. 2005;41:2449–2461. doi: 10.1016/j.ejca.2005.08.008

39. Hu X, Li YQ, Li QG, Ma YL, Peng JJ, Cai SJ. Osteoglycin (OGN) reverses epithelial to mesenchymal transition and invasiveness in colorectal cancer via EGFR/Akt pathway. J Exp Clin Cancer Res. 2018;37:41. doi: 10.1186/s13046-018-0718-2

40. Angiolilli C, Leijten EFA, Bekker CPJ, Eeftink E, Giovannone B, Nordkamp MO, van der Wal M, Thijs JL, Vastert SJ, van Wijk F, et al. ZFP36 Family Members Regulate the Proinflammatory Features of Psoriatic Dermal Fibroblasts. J Invest Dermatol. 2022;142:402–413. doi: 10.1016/j.jid.2021.06.030

41. Kobayashi A, Takahashi T, Horita S, Yamamoto I, Yamamoto H, Teraoka S, Tanabe K, Hosoya T, Yamaguchi Y. Activation of the transcription factor c-Jun in acute cellular and antibody-mediated rejection after kidney transplantation. Hum Pathol. 2010;41:1682–1693. doi: 10.1016/j.humpath.2010.04.016

42. Mintet E, Lavigne J, Paget V, Tarlet G, Buard V, Guipaud O, Sabourin JC, Iruela-Arispe ML, Milliat F, François A. Endothelial Hey2 deletion reduces endothelial-to-mesenchymal transition and mitigates radiation proctitis in mice. Sci Rep. 2017;7:4933. doi: 10.1038/s41598-017-05389-8

43. Wang XM, Liu XM, Wang Y, Chen ZY. Activating transcription factor 3 (ATF3) regulates cell growth, apoptosis, invasion and collagen synthesis in keloid fibroblast through transforming growth factor beta (TGF-beta)/SMAD signaling pathway. Bioengineered. 2021;12:117–126. doi: 10.1080/21655979.2020.1860491

44. Malabanan KP, Kanellakis P, Bobik A, Khachigian LM. Activation transcription factor-4 induced by fibroblast growth factor-2 regulates vascular endothelial growth factor-A transcription in vascular smooth muscle cells and mediates intimal thickening in rat arteries following balloon injury. Circ Res. 2008;103:378–387. doi: 10.1161/circresaha.107.168682

45. Chen X, Xu J, Bao W, Li H, Wu W, Liu J, Pi J, Tomlinson B, Chan P, Ruan C, et al. Endothelial Foxp1 Regulates Neointimal Hyperplasia Via Matrix Metalloproteinase-9/Cyclin Dependent Kinase Inhibitor 1B Signal Pathway. J Am Heart Assoc. 2022;11:e026378. doi: 10.1161/jaha.122.026378

46. Whitmore HAB, Amarnani D, O’Hare M, Delgado-Tirado S, Gonzalez-Buendia L, An M, Pedron J, Bushweller JH, Arboleda-Velasquez JF, Kim LA. TNF-α signaling regulates RUNX1 function in endothelial cells. Faseb j. 2021;35:e21155. doi: 10.1096/fj.202001668R

47. Berry GJ, Burke MM, Andersen C, Bruneval P, Fedrigo M, Fishbein MC, Goddard M, Hammond EH, Leone O, Marboe C, et al. The 2013 International Society for Heart and Lung Transplantation Working Formulation for the standardization of nomenclature in the pathologic diagnosis of antibody-mediated rejection in heart transplantation. J Heart Lung Transplant. 2013;32:1147–1162. doi: 10.1016/j.healun.2013.08.011

48. Lu WH, Palatnik K, Fishbein GA, Lai C, Levi DS, Perens G, Alejos J, Kobashigawa J, Fishbein MC. Diverse morphologic manifestations of cardiac allograft vasculopathy: a pathologic study of 64 allograft hearts. J Heart Lung Transplant. 2011;30:1044–1050. doi: 10.1016/j.healun.2011.04.008

49. Valenzuela NM, Hong L, Shen XD, Gao F, Young SH, Rozengurt E, Kupiec-Weglinski JW, Fishbein MC, Reed EF. Blockade of p-selectin is sufficient to reduce MHC I antibody-elicited monocyte recruitment in vitro and in vivo. Am J Transplant. 2013;13:299–311. doi: 10.1111/ajt.12016

50. Wei X, Valenzuela NM, Rossetti M, Sosa RA, Nevarez-Mejia J, Fishbein GA, Mulder A, Dhar J, Keslar KS, Baldwin WM, 3rd, et al. Antibody-induced vascular inflammation skews infiltrating macrophages to a novel remodeling phenotype in a model of transplant rejection. Am J Transplant. 2020;20:2686–2702. doi: 10.1111/ajt.15934

51. Choy JC, Cruz RP, Kerjner A, Geisbrecht J, Sawchuk T, Fraser SA, Hudig D, Bleackley RC, Jirik FR, McManus BM, et al. Granzyme B induces endothelial cell apoptosis and contributes to the development of transplant vascular disease. Am J Transplant. 2005;5:494–499. doi: 10.1111/j.1600-6143.2004.00710.x

52. Choy JC, McDonald PC, Suarez AC, Hung VH, Wilson JE, McManus BM, Granville DJ. Granzyme B in atherosclerosis and transplant vascular disease: association with cell death and atherosclerotic disease severity. Mod Pathol. 2003;16:460–470. doi: 10.1097/01.Mp.0000067424.12280.Bc

53. Adachi M, Imai K. The proapoptotic BH3-only protein BAD transduces cell death signals independently of its interaction with Bcl-2. Cell Death & Differentiation. 2002;9:1240–1247. doi: 10.1038/sj.cdd.4401097

54. Jin YP, Fishbein MC, Said JW, Jindra PT, Rajalingam R, Rozengurt E, Reed EF. Anti-HLA class I antibody-mediated activation of the PI3K/Akt signaling pathway and induction of Bcl-2 and Bcl-xL expression in endothelial cells. Hum Immunol. 2004;65:291–302. doi: 10.1016/j.humimm.2004.01.002

55. Chatterjee D, Moore C, Gao B, Clerkin KJ, See SB, Shaked D, Rogers K, Nunez S, Veras Y, Addonizio L, et al. Prevalence of polyreactive innate clones among graft--infiltrating B cells in human cardiac allograft vasculopathy. J Heart Lung Transplant. 2018;37:385–393. doi: 10.1016/j.healun.2017.09.011

56. Wehner J, Morrell CN, Reynolds T, Rodriguez ER, Baldwin WM, 3rd. Antibody and complement in transplant vasculopathy. Circ Res. 2007;100:191–203. doi: 10.1161/01.Res.0000255032.33661.88

57. Tsuda H, Dvorina N, Keslar KS, Nevarez-Mejia J, Valenzuela NM, Reed EF, Fairchild RL, Baldwin WM, 3rd. Molecular Signature of Antibody-Mediated Chronic Vasculopathy in Heart Allografts in a Novel Mouse Model. Am J Pathol. 2022;192:1053–1065. doi: 10.1016/j.ajpath.2022.04.003

58. Huibers MM, Vink A, Kaldeway J, Huisman A, Timmermans K, Leenders M, Schipper ME, Lahpor JR, Kirkels HJ, Klöpping C, et al. Distinct phenotypes of cardiac allograft vasculopathy after heart transplantation: a histopathological study. Atherosclerosis. 2014;236:353–359. doi: 10.1016/j.atherosclerosis.2014.07.016

59. Hruban RH, Beschorner WE, Baumgartner WA, Augustine SM, Ren H, Reitz BA, Hutchins GM. Accelerated arteriosclerosis in heart transplant recipients is associated with a T-lymphocyte-mediated endothelialitis. Am J Pathol. 1990;137:871–882.

60. Fu Y, Lin Q, Zhang Z, Zhang L. Therapeutic strategies for the costimulatory molecule OX40 in T-cell-mediated immunity. Acta Pharm Sin B. 2020;10:414–433. doi: 10.1016/j.apsb.2019.08.010

61. Tkachev V, Furlan SN, Watkins B, Hunt DJ, Zheng HB, Panoskaltsis-Mortari A, Betz K, Brown M, Schell JB, Zeleski K, et al. Combined OX40L and mTOR blockade controls effector T cell activation while preserving T(reg) reconstitution after transplant. Sci Transl Med. 2017;9. doi: 10.1126/scitranslmed.aan3085

62. Morrell CN, Murata K, Swaim AM, Mason E, Martin TV, Thompson LE, Ballard M, Fox-Talbot K, Wasowska B, Baldwin WM, 3rd. In vivo platelet-endothelial cell interactions in response to major histocompatibility complex alloantibody. Circ Res. 2008;102:777–785. doi: 10.1161/circresaha.107.170332

63. Valenzuela NM, Mulder A, Reed EF. HLA class I antibodies trigger increased adherence of monocytes to endothelial cells by eliciting an increase in endothelial P-selectin and, depending on subclass, by engaging FcγRs. J Immunol. 2013;190:6635–6650. doi: 10.4049/jimmunol.1201434

64. He L, Li Q, Du C, Xue Y, Yu P. Glis2 inhibits the epithelial-mesenchymal transition and apoptosis of renal tubule cells by regulating the β-catenin signalling pathway in diabetic kidney disease. Biochem Biophys Res Commun. 2022;607:73–80. doi: 10.1016/j.bbrc.2022.03.111

65. Jin YP, Valenzuela NM, Ziegler ME, Rozengurt E, Reed EF. Everolimus inhibits anti-HLA I antibody-mediated endothelial cell signaling, migration and proliferation more potently than sirolimus. Am J Transplant. 2014;14:806–819. doi: 10.1111/ajt.12669

66. Jin YP, Singh RP, Du ZY, Rajasekaran AK, Rozengurt E, Reed EF. Ligation of HLA class I molecules on endothelial cells induces phosphorylation of Src, paxillin, and focal adhesion kinase in an actin-dependent manner. J Immunol. 2002;168:5415–5423. doi: 10.4049/jimmunol.168.11.5415

67. Tavora F, Munivenkatappa R, Papadimitriou J, Drachenberg C, Sailey C, Mehra M, Burke A. Endothelitis in cardiac allograft biopsy specimens: possible relationship to antibody-mediated rejection. J Heart Lung Transplant. 2011;30:435–444. doi: 10.1016/j.healun.2010.10.009

